# SquiDBase: a community resource of raw nanopore data from microbes

**DOI:** 10.1101/2025.04.28.650941

**Authors:** Wim L. Cuypers, Halil Ceylan, Eline Turcksin, Laura Raes, Nicky de Vrij, Johan Michiels, Sandra Coppens, Tessa de Block, Daan Jansen, Kevin K. Ariën, Philippe Selhorst, Koen Vercauteren, Julia M. Gauglitz, Wout Bittremieux, Kris Laukens

## Abstract

Experimental data-driven research relies on raw data, which consist of unprocessed experimental outputs, whereas derived data are transformed through a number of processing steps to reveal specific insights. Such processing, however, can potentially introduce biases or information loss, compromising transparency and reproducibility. In nucleic acid sequencing, nucleotide sequences stored in the FASTQ format are widely shared, but FASTQ files are generated from platform-specific raw data outputs, which vary depending on the sequencing platform used. The raw data produced by Oxford Nanopore Technologies (ONT) sequencing devices contain valuable biological information and are also useful to improve data processing methods, which includes basecaller optimisation and modification detection. Increasing attention goes to exploring these raw signals to develop algorithms that could improve ONT device portability and enhance target enrichment efficiency through adaptive sampling. Despite these benefits, the storage and sharing of raw nanopore data remain limited due to technical constraints and the lack of appropriate, standardised and centralised infrastructure. To address this challenge, we developed SquiDBase (https://squidbase.org), a dedicated repository to collect raw microbial nanopore sequencing data. To maximise the utility of SquiDBase from its inception, we built SquiDPipe, a Nextflow pipeline for the automated removal of human or unwanted reads from raw nanopore data. Additionally, we sequenced 24 clinically relevant viruses and incorporated them into SquiDBase, significantly expanding the diversity of publicly available reference datasets. By offering a centralised, open-access raw data collection platform, SquiDBase facilitates data sharing, enhances reproducibility, and supports the development and benchmarking of novel computational tools, reinforcing open science in nanopore sequencing research.

## INTRODUCTION

In this era of data-intensive experimental science and open data policies, raw data—the unmodified output of experimental instruments—serves as the foundational record of scientific observations. In contrast, derived data are processed or analysed versions of raw data, tailored to generate answers to specific research questions^1^, potentially introducing biases or information loss. Sharing raw data with sufficient metadata enhances reproducibility, enables meta-analyses, and supports new analytical methods^2–4^. In the nucleic acid sequencing field, the FASTQ file format is the standard for representing raw sequencing data in a uniform and portable manner. Public repositories such as the European Nucleotide Archive (ENA)^5^ and the Sequence Read Archive (SRA)^6^ have facilitated the widespread sharing of these data originating from different sequencing technologies. While this practice has greatly facilitated data sharing, it is worth considering whether sharing even rawer forms of sequencing data—those generated directly by the sequencing instruments—could further advance open science and improve reproducibility.

Sequencing reads in FASTQ files contain both nucleotide sequences and their associated quality scores^7^, generated through a process known as ‘basecalling’^8^. The nature of the initial raw data, its utility, and the need for long-term storage, varies significantly across sequencing platforms. In fluorescence-based sequencing-by-synthesis (SBS) technologies, such as those used by Illumina and PacBio, nucleotide incorporation signals are captured and automatically processed into high-quality FASTQ files^9^, reducing the need for user interaction with raw data or its storage. While these methods provide highly accurate nucleotide sequences, extracting additional biological insights such as DNA methylation requires specialised preprocessing (*e*.*g*., bisulphite conversion) or data beyond signal intensities, as seen in PacBio’s HiFi sequencing, which infers epigenetic information from polymerase kinetics at the single-molecule level^10^.

In contrast, nanopore sequencing often necessitates direct engagement with raw signal data. Nanopore sequencers detect changes in ionic current in real-time as nucleic acids pass through a nanopore, enabling the detection of their native state^11,12^. This creates multiple incentives for retaining and analysing raw nanopore data. First, basecaller accuracy on native DNA improves when training data includes species-specific modifications, as these alter raw signal patterns^13–15^. Second, raw data support downstream analyses, including the detection of methylation^16,17^ and other modifications^18^, splice-junction identification^19^, and the characterisation of tandem repeats^20–22^. Third, advancements in real-time analysis, such as live basecalling and adaptive sampling for target enrichment, have intensified the search for efficient raw signal processing. While graphics processing unit (GPU)-based basecalling remains computationally demanding, alternative approaches are emerging capable of real-time analysis during portable sequencing using minimal hardware^23,24^ and the simultaneous processing of data from multiple sequencing devices^25^. These methods include UNCALLED^26^, RawAlign/RawHash2^27,28^, Sigmoni^29^, SigMap^30^, BaseLess^25^, and SquiggleNet^31^. For all these applications, development and benchmarking remain constrained by the limited availability of high-quality raw data from diverse species.

Despite the clear benefits of preserving raw nanopore data for experimental researchers and algorithm developers, enabling re-analysis, meta-analyses, and benchmarking, it is rarely shared in the public domain^32,33^. We hypothesise that this is due to both storage limitations and the lack of support for raw nanopore data formats. Consequently, many users upload only processed data in the FASTQ format, yet the information within these files becomes outdated with each new release of basecalling models, leading to data loss.

To overcome these challenges and ensure long-term accessibility of raw nanopore data in accordance with open-science principles, we developed SquiDBase (short for “Squiggle database”), a repository designed for storing and sharing raw nanopore signals from microbial sequencing projects (encompassing prokaryotes, microbial eukaryotes, and viruses). Our focus on microbial data stems from their scalability, reduced privacy constraints compared to human genomics, and their critical role in pandemic preparedness. This emphasis also fills an important gap, as current large-scale nanopore sequencing efforts prioritise human genomics, leaving microbial datasets, for instance those needed for benchmarking taxonomic classifiers^34^, relatively scarce. By centralising these data, SquiDBase significantly enhances the accessibility and reusability of raw microbial nanopore data, facilitating experimental research and the development and benchmarking of new algorithms.

## MATERIALS AND METHODS

SquiDBase is a specialised repository designed to store raw nanopore data in the POD5 file format, along with associated metadata. SquiDBase is particularly tailored for data originating from microbial species, making it a valuable resource for researchers studying infectious diseases and metagenomics of humans and the environment. The database is accessible online via https://SquiDBase.org, offering researchers a user-friendly platform to upload, store, and retrieve nanopore sequencing data and metadata.

### Database structure and implementation

SquiDBase stores all metadata linked to a set of raw nanopore reads in a PostgreSQL database, structured to adhere to Third Normal Form (3NF). This design minimizes redundancy and ensures data integrity by organizing tables so that all non-key attributes depend solely on primary keys. We employ Alembic for database migrations, enabling us to track changes and maintain version control over the database schema. This ensures consistent application of updates and modifications across environments.

In total, 11 tables contain all information linked to a given submission in SquiDBase (**Supplementary Table 1; Supplementary Figure 1**). These include tables for user data (*users*), technical details associated with the nanopore sequencing run (*pod5_read, nanopore_kit*), metadata related to the sample that was sequenced (*pod5_file, country, source, ncbi_taxon, diagnostic*), and a table for managing submissions (*submissions*).

### Data upload and standardisation

To contribute to SquiDBase, users must create an account before they can upload data. Facilitating a secure, efficient, and responsive user experience, SquiDBase incorporates a registration and authentication system backend powered by FastAPI, while the frontend leverages Nuxt.js. User sessions are maintained using JSON Web Tokens (JWTs), which securely transmit information between parties and authenticate users, keeping their sessions active without compromising security.

Once registered, raw nanopore data in the POD5 format can be uploaded along with comprehensive metadata in a predefined CSV format to enhance the utility of the dataset. Metadata fields include filename, the taxonomic identifier of the predominant species in the sample, and additional metadata fields detailed in **Table 1**. Where applicable, we enhance standardisation by using established ontologies and controlled vocabularies. For example, we employ UBERON^35^, ENVO^36^, and FOODON^37^ to specify the sample source, and the OBI ontology^38^ to describe the diagnostic method. Additionally, we use NCBI Taxonomy^39^ to classify biological organisms and ISO 3166 two-letter country codes to indicate the country of origin and isolation (**Table 1**).

**Table 1:** Metadata fields in SquiDBase accompanying each upload. Standardised ontologies and conventions were used where possible, such as UBERON for source classification and ISO 3166 for two-letter country codes. The table includes the field name, a description, the data type, and an example for each metadata field.

| Field/Variable name | Description | Data type | Example |
| --- | --- | --- | --- |
| **Filename** | The file name of the POD5 sequencing data to be uploaded. | String | 37124_CHIKV-1.pod5 |
| **Species TaxID** | NCBI taxonomy identifier for the microbial species in the sample. | NCBI taxonomy | 37124 |
| **Year of isolation** | The year the pathogen was collected or isolated from the host. For non-pathogenic organisms and metagenomic samples, this refers to the year the sample was collected. | Integer | 2014 |
| **Country of isolation** | The country where the organism was isolated or the sample was collected. | ISO 3166 country code | BE |
| **Geographic origin** | The likely country of origin of the organism or infection, if known. This may differ from the country of isolation—for example, in travel-related infections. For pathogens, this refers to the most probable geographic source of the infection. | ISO 3166 country code | ET |
| **Strain or lineage** | The specific strain, lineage, or sequence type of the uploaded pathogen data. | String | BA.5 |
| **Source ID** | A unique identifier for the source of the pathogen sample, utilizing the UBERON, ENVO, or FOODON ontologies. | UBERON, ENVO, and FOODON ontologies | UBERON:0000178 |
| **Host TaxID** | NCBI taxonomy identifier for the host species from which the pathogen was isolated, if applicable. | NCBI taxonomy | 9606 |
| **Internal lab ID** | The internal identification code assigned to the sample by the laboratory. | String | PLAS-ETH-2023-0147 |
| **Diagnostic method** | The diagnostic method used to detect the pathogen, if applicable, is represented using the OBI ontology. | OBI ontology | OBI:0003045 |
| **Remarks** | Additional notes or remarks about the sample, often for internal collection records. | Text field | “Strain donated by institute X.”, or “Collected for the SquiDBase project”. |

Each dataset includes an information field that supports free-text input, allowing users to provide essential details about the data. This field is particularly valuable for documenting the rationale and methodology of their study, with a specific emphasis on the wet-lab procedures used to generate the data. For instance, if PCR amplification was employed to increase input material, or if similar techniques were used that remove or alter base modifications, these steps should be explicitly noted. Such details are important, as preprocessing methods can significantly influence the raw signal data and its subsequent interpretation. Providing this information increases transparency and improves the accuracy of downstream analyses.

During upload, SquiDBase automatically extracts and stores comprehensive sequencing run metadata from submitted POD5 files, including the type of experiment, device model, flow cell specifications, used library preparation kit, acquisition parameters, and real-time throughput (**Table 2**).

**Table 2:** Information on the sequencing run that is extracted from POD5 files and stored in a separate ‘pod5_file’ table and other tables as needed to maintain third normal form (3NF) within SquiDBase.

| Field from POD5 file | Description | Example |
| --- | --- | --- |
| **context_tags.experiment_type** | Type of experiment conducted. | genomic_dna |
| **sample_rate** | Sequencing run sample rate in samples per second. | 4000 |
| **context_tags.selected_speed_bases_per_second** | Approximate number of bases read per second during sequencing. | 400 |
| **sequencer_position_type** | Model of sequencing device used. | MinION Mk1B |
| **sequencing_kit** | Sequencing kit used for library preparation. | sqk-lsk114 |
| **flow_cell_product_code** | Product code indicating the flow cell type. | FLO-MIN114 |

### Preprocessing pipeline for human read removal or species extraction

By default, SquiDBase assumes that the metadata associated with a POD5 file apply to all reads within that file. This assumption typically holds true for pure samples, where all reads originate from the same source. However, in cases where the sample is mixed, such as a patient sample containing both human and viral data, the viral raw signals must be extracted before uploading to SquiDBase, as the platform is designed for sharing microbial, not human data. To facilitate this process, we have developed SquiDPipe (https://github.com/SquiDBase/SquiDPipe), a Nextflow-based pipeline that segments and filters POD5 files to isolate unaltered signals of interest.

Users initiate the pipeline by providing the paths to demultiplexed FASTQ and POD5 files, which are organized by barcode (*e*.*g*., each barcode folder contains all files for that barcode). Alternatively, users can provide a single folder containing all POD5 files, though this increases processing time. Additionally, a CSV file must be supplied, mapping each barcoded FASTQ folder to the species name and corresponding NCBI taxonomic identifier for targeted read extraction. This ensures the pipeline can accurately identify and isolate reads of interest for further processing.

The pipeline first processes the basecalled FASTQ files to detect reads originating from species of interest. It performs taxonomic read classification on all reads within each barcode, recording the results. For optimal accuracy, a targeted, organism-specific database (*e*.*g*., RefSeq virus) needs to be used.

Once reads associated with target organisms are identified, the pipeline retrieves reference genomes from RefSeq. If a reference genome is unavailable in RefSeq for a specific taxonomic identifier, the pipeline defaults to GenBank. In cases where neither RefSeq nor GenBank contain a genome, the pipeline attempts to locate a reference genome for the parent taxon using NCBI taxonomy information. These reference genomes, along with any additional user-specified genomes (*e*.*g*., the human reference genome), are compiled into a single FASTA file. To reduce false positive identifications, the pipeline maps the identified reads to the combined reference genome. Reads that accurately map to the reference genome of the correct species are extracted, and their read IDs are used to filter the corresponding POD5 files. This process generates POD5 files containing only the reads from the target species.

The final output consists of POD5 files containing reads from a single target species, ready for upload to SquiDBase, along with mapping statistics for all species that were extracted. If complete metadata were provided as input, SquiDPipe additionally generates a CSV file formatted for upload to SquiDBase, ensuring seamless integration with the platform.

### Data retrieval and access

Once uploaded, raw nanopore data are accessible to the scientific community for reanalysis and further research. Users can search for datasets via the ‘Browse Datasets’ tab (accessible at https://squidbase.org/submissions) (**Supplementary Figure 2**). To facilitate user exploration, the page includes a filtering feature that allows users to filter datasets based on the nanopore flow cell type (*e*.*g*., R9.4.1 or R10.4.1). This functionality is particularly valuable as the flow cell type is a critical factor influencing raw signal characteristics, which is essential for developing and optimizing analytical algorithms.

Users can access each dataset through a unique SquiDBase identifier (*e*.*g*. https://squidbase.org/submissions/SQB000004). Each dataset page includes a brief text summary, a visual overview of the metadata, and a section for downloading the data directly from the browser (**Supplementary Figure 3**). A query functionality further enables users to search for files based on specific metadata (*e*.*g*., taxonomic identifier) within a submission. Users can access full metadata for any file by clicking an eye icon, which also provides an MD5 checksum for file integrity verification (**Supplementary Figure 4**).

SquiDBase employs presigned URLs for secure file transfers, which expire after 24h to prevent unauthorized access or hotlinking. Users access files via the submission page, maintaining dataset context. JSON-RPC facilitates smooth communication between the Nuxt.js frontend and the FastAPI backend, ensuring efficient data exchange for user authentication and core functionalities.

Expansion and maintenance of the system are further facilitated by FastAPI’s built-in support for OpenAPI, which provides comprehensive API documentation. This feature simplifies the integration of new frontends, such as Python libraries or command-line tools, in the future. Developers can easily access and understand the available endpoints, ensuring that the system remains adaptable and scalable as new features or tools are introduced.

### Legacy R9 data and expanding public R10.4.1 collections

To support benchmarking, we centralised legacy datasets generated using the older R9 flowcells from sources like SRA and CADDE on AWS into SquiDBase. These datasets have been previously used for benchmarking tools such as RawHash, Sigmoni and others^26,28,29,31^. The collection includes R9 datasets for *E. coli*, SARS-CoV-2, and *Chlamydomonas reinhardtii*, and were converted from FAST5 to POD5 using the POD5 package v0.3.15^40^ and merged into larger files.

Publicly available R10.4.1 sequencing data for microbes remain limited and scattered. To address this gap, and given the significance of viruses in pandemic preparedness, we prioritised sequencing clinically 20 relevant viral species (68 isolates). Included in SquiDBase were Chikungunya virus (CHIKV-1, CHIKV-2, CHIKV-3, CHIKV-4), Eastern Equine Encephalitis virus (EEEV), Dengue virus (DENV1, DENV2, DENV3, DENV4), Human Immunodeficiency virus (HIV), Japanese Encephalitis virus (JEV), Mayaro virus (MAYV), Monkeypox virus (MPOX), O’nyong-nyong virus (ONNV), Rift Valley Fever virus (RVFV), Sandfly Fever Naples virus (SFNV), Sandfly Fever Turkey virus (SFTV), Sindbis virus (SINV), Severe Acute Respiratory Syndrome Coronavirus 2 (SARS-CoV-2), Tick-borne Encephalitis virus (TBEV), Usutu virus (USUV), Venezuelan equine encephalitis virus (VEEV), West Nile virus (WNV), Western Equine Encephalitis virus (WEEV), Yellow Fever virus (YFV), and Zika virus (ZIKV). In addition, we added data originating from *Plasmodium falciparum*, which is responsible for the majority of malaria cases and frequently harbours drug resistance mutations. Wet-lab procedures, sequencing and bioinformatics processing are described in **Supplementary Methods**.

## RESULTS AND DISCUSSION

We developed SquiDBase to promote the sharing of raw nanopore data in POD5 format originating from microbial sources by ensuring they are findable, accessible, interoperable, and reusable (FAIR).

SquiDBase provides a streamlined process for researchers to share and access raw nanopore data, even without prior bioinformatics expertise. Data can be uploaded using a unique SquiDBase identifier, allowing the community to build upon existing data generation efforts and maintain consistent benchmarks over time. Submission methods vary by sample type: single-specimen or single-species datasets can be uploaded directly, while clinical samples containing human and microbial reads require preprocessing with the SquiDPipe pipeline which we developed to remove human data. Metagenomic samples composed of multiple non-human species must include the appropriate NCBI taxonomic identifier for accurate classification. Once deposited, datasets remain openly accessible, with users able to retrieve raw nanopore data in POD5 format via the browse page or direct submission URL (*e*.*g*. https://squidbase.org/submissions/SQB000004), without login requirements. Comprehensive documentation and preprocessing tools further enhance the utility of SquiDBase, making it a robust resource for the research community.

To maintain compatibility with prior benchmarking efforts, we have integrated datasets generated with R9 flow cell types, which were previously dispersed across multiple sources. These datasets are now centralised and accessible from the dataset overview page. Given the limited availability of recent R10.4.1 data, we have prioritised expanding the collection of publicly available raw nanopore data. This focused on viral datasets, contributing data for 68 viral isolates, encompassing over 26 clinically relevant viruses from multiple species and serotypes. Descriptions on sequencing depth and coverage for each of the viral isolates sequenced for this project can be found in **Supplementary Table 2**. Furthermore, we have contributed raw sequencing data of *Plasmodium falciparum*, the predominant malaria-causing species. Data from four samples derived from three human patients were included. Chromosome-wide coverage statistics for these *P. falciparum* samples are provided in **Supplementary Table 3**. Overall this will serve as a foundation for developing new algorithms such as species-specific basecallers and advancing targeted research efforts.

Looking ahead, our roadmap prioritises expanding microbial R10.4.1 datasets in SquiDBase while improving querying capabilities at the POD5 read level. Key future developments include enhanced metadata visualization, raw signal matching, and alignment features to facilitate modification analysis and rapid signal-based searches. As SquiDBase evolves, it will become a more comprehensive platform for storing, retrieving, and analysing raw nanopore data from microbes. Managing the large file sizes associated with raw nanopore data remains a major challenge. While alternative formats such as SLOW5 exist, we continue to use POD5, the format endorsed by ONT, despite its trade-offs. At present, our focus remains on microbial datasets due to their relatively small genome size, but future developments—driven by technological advancements and community demand—may allow us to broaden the scope to additional organisms and explore more efficient file formats.

In conclusion, SquiDBase addresses the urgent need for a centralised repository of raw nanopore data, promoting open science and enabling the research community to fully leverage nanopore sequencing technology. Worth mentioning is the analogy to the field of proteomics, where the practice of sharing raw mass spectrometry data in public repositories has driven algorithm improvements and novel discoveries^41^. Following this model, we propose that SquiDBase will improve the long-term research utility of nanopore sequencing data, catalysing breakthroughs in diagnostics, base modification research, and the development of algorithms that can process vast amounts of nanopore data in resource-limited settings. Central to these innovations is the availability of extensive datasets for model training, re-analysis, meta-analyses, and benchmarking.

## FUNDING & ACKNOWLEDGEMENTS

This work was supported by the Industrial Research Fund of the University of Antwerp and ITM’s SOFI programme supported by the Flemish Government, Science & Innovation. In addition, WLC is supported by a Flanders Innovation & Entrepreneurship (VLAIO) Innovation Mandate (HBC.2024.0273).

