## Supplementary materials for "SquiDBase: a community resource of raw nanopore data from microbes"

**Supplementary Methods**

Virus samples were processed by the Clinical Virology and Virology Units at the Institute of Tropical Medicine Antwerp (ITM). For monkeypox virus, DNA was extracted and amplified using sequence-independent single-primer amplification (SISPA), following the protocol described by De Baetselier et al. (2022)[1]. For RNA viruses, the starting material consisted of culture supernatant obtained from the Virology Unit virus repository. The extraction and enrichment protocol included RNA extraction, TurboDNase treatment, reverse transcription to cDNA, and subsequent SISPA amplification, in line with previous publications by Greninger et al. (2015)[2] and Kafetzopoulou et al. (2018)[3]. Sequencing of the viral strains was carried out at the University of Antwerp using the ONT MinION platform on R10.4.1 flow cells (FLO-MIN114). Library preparation was performed using the Rapid Barcoding Kit v14 (SQK-RBK114.24), and barcoded libraries were sequenced in batches of 24 pooled samples per flow cell for 42 to 72 hours. The output ranged from 1.88 up to 5.22 gigabase. Raw nanopore reads in POD5 format were basecalled using Dorado (ONT Dorado basecall server software version 7.0.2+7e7b7d0) employing the super-accurate (SUP) model (dna_r10.4.1_e8.2_400bps_5khz). Data were further processed using SquiDPipe v1.0.0 (https://github.com/Cuypers-Wim/squidpipe), implemented in Nextflow v24.04.4[4]. In brief, reads were classified and extracted using Kraken2 v2.1.3[5] with the RefSeq viral database[6] (downloaded May 25, 2024) and Seqtk v1.4-r122[7]. Reference genomes for target species and human (to account for contamination) were retrieved using the NCBI datasets CLI v16.27.1[8]. Reads were aligned to concatenated reference genomes using Minimap2 v2.28-r1209[9], and alignments were processed with Samtools v1.20[10,11] to remove duplicates, generate BAM files, and calculate coverage statistics. Finally, the Pod5 package v0.3.15[12] was used to extract reads mapping exclusively to the target species, consolidating them into single POD5 files per species.

To further expand the pathogen diversity represented in SquiDBase, we incorporated raw nanopore whole-genome sequencing data from Plasmodium falciparum, a clinically significant pathogen for which such data were previously unavailable. The complete wet lab protocol is available in the study by De Meulenaere et al. (2024)[13]. We ran SquiDPipe v1.0.0 to extract raw nanopore signal data specific to Plasmodium. Using this approach, we obtained data for Plasmodium falciparum from three human patients, with native parasite DNA sequenced directly from blood using nanopore sequencing in adaptive sampling mode. For one of these samples, we also generated sWGA-enriched data in addition to the native DNA dataset.

### Supplementary Tables

***Supplementary Table 1**: Description of the PostgreSQL database tables in SquiDBase.*

| Table name | Description |
|---|---|
| **users** | Stores user information. Essential for managing user accounts and linking them to submissions. |
| **country** | Holds country data using ISO codes. Used to reference geographic locations in other tables. |
| **nanopore_chemistry** | Lists types of nanopore chemistries. |
| **nanopore_kit** | Contains details about various nanopore kits. |
| **source** | Stores source information from various ontologies (UBERON, FOODON, ENVO). Provides standardized references. |
| **submissions** | Manages data submission records. |
| **ncbi_taxon** | Contains taxonomic data with hierarchical relationships, essential for biological classification. |
| **pod5_file** | Represents files with metadata linking to countries, taxonomies, and submissions for comprehensive tracking. |
| **pod5_read** | Stores read-level data linked to files. Ensures each read is associated with a valid file. |
| **diagnostic** | Stores source information from OBI ontology. Provides standardized references. |
| **alembic_version** | Maintains a record of applied database schema migrations, ensuring version control and reproducibility across development environments. It provides standardized tracking of structural changes to the database. |

***Supplementary Table 2****: Coverage statistics for virus isolates in SquiDBase. The table includes the following columns: "Name" (virus abbreviation, e.g., DENV1), "StudyID" (internal lab identifier), "NCBI TaxID" (taxonomic identifier), "Chromosome" (RefSeq identifier of the reference genome used for read mapping), "Coverage" (proportion of the reference genome covered by at least one read), "Depth of Coverage" (average number of reads mapped to a position in the reference genome). The remaining columns indicate the percentage of positions in the reference genome covered by at least 10, 20, or 30 reads respectively.*

| Name | studyID | NCBI TaxID | Chromosome | Coverage (%) | Depth of Coverage | Positions > 10 reads (%) | Positions > 20 reads (%) | Positions > 30 reads (%) |
|---|---|---|---|---|---|---|---|---|
| **CHIKV** | 1 | 37124 | NC_004162.2 | 100.0 | 486.1 | 98.4 | 97.6 | 96.6 |
| **CHIKV** | 2 | 37124 | NC_004162.2 | 100.0 | 739.9 | 99.7 | 99.1 | 98.6 |
| **CHIKV** | 3 | 37124 | NC_004162.2 | 99.87 | 299.8 | 99.1 | 98.5 | 97.6 |
| **CHIKV** | 4 | 37124 | NC_004162.2 | 99.94 | 75.3 | 82.3 | 76.3 | 70.3 |
| **DENV1** | 5 | 11053 | NC_001477.1 | 99.96 | 602.5 | 98.9 | 98.3 | 98.2 |
| **DENV1** | 6 | 11053 | NC_001477.1 | 99.59 | 764.8 | 99.0 | 98.9 | 98.8 |
| **DENV2** | 7 | 11060 | NC_001474.2 | 99.68 | 531.7 | 98.8 | 98.3 | 97.9 |
| **DENV2** | 8 | 11060 | NC_001474.2 | 99.67 | 973.4 | 99.1 | 99.1 | 98.9 |
| **DENV2** | 9 | 11060 | NC_001474.2 | 99.42 | 208.3 | 98.2 | 96.6 | 96.1 |
| **DENV3** | 10 | 11069 | NC_001475.2 | 99.05 | 232.0 | 98.8 | 98.0 | 97.6 |
| **DENV3** | 11 | 11069 | NC_001475.2 | 99.97 | 772.4 | 99.1 | 98.9 | 98.8 |
| **DENV3** | 12 | 11069 | NC_001475.2 | 99.84 | 172.8 | 98.3 | 97.3 | 95.9 |
| **DENV3** | 68 | 11069 | NC_001475.2 | 99.69 | 685.0 | 99.1 | 99.1 | 98.9 |
| **DENV4** | 13 | 11070 | NC_002640.1 | 99.97 | 667.4 | 99.4 | 99.1 | 98.7 |
| **DENV4** | 14 | 11070 | NC_002640.1 | 99.94 | 173.8 | 98.1 | 97.4 | 96.9 |
| **DENV4** | 15 | 11070 | NC_002640.1 | 99.68 | 227.3 | 98.5 | 97.5 | 97.4 |
| **EEEV** | 16 | 11021 | NC_003899.1 | 99.99 | 544.5 | 99.7 | 99.6 | 99.5 |
| **HIV** | 59 | 11676 | NC_001802.1 | 100.0 | 332.2 | 97.2 | 96.7 | 96.2 |
| **HIV** | 60 | 11676 | NC_001802.1 | 99.93 | 299.5 | 96.7 | 93.7 | 92.3 |
| **HIV** | 61 | 11676 | NC_001802.1 | 99.93 | 552.7 | 96.9 | 95.2 | 93.8 |
| **HIV** | 62 | 11676 | NC_001802.1 | 99.98 | 414.3 | 97.4 | 96.4 | 95.7 |
| **HIV** | 63 | 11676 | NC_001802.1 | 99.84 | 221.6 | 96.2 | 93.4 | 90.3 |
| **HIV** | 64 | 11676 | NC_001802.1 | 97.79 | 193.00 | 93.1 | 88.1 | 85.7 |
| **JEV** | 17 | 11072 | NC_001437.1 | 99.99 | 508.8 | 99.1 | 99.0 | 99.0 |
| **JEV** | 18 | 11072 | NC_001437.1 | 100.0 | 763.8 | 99.3 | 99.1 | 99.0 |
| **JEV** | 19 | 11072 | NC_001437.1 | 99.45 | 198.5 | 99.0 | 98.9 | 98.8 |
| **JEV** | 20 | 11072 | NC_001437.1 | 99.13 | 199.5 | 99.0 | 98.8 | 98.8 |
| **MAYV** | 21 | 59301 | NC_003417.1 | 98.09 | 215.8 | 97.8 | 97.6 | 94.5 |
| **MPOX** | 65 | 10244 | NC_063383.1 | 45.67 | 0.7 | 0 | 0 | 0 |
| **MPOX** | 66 | 10244 | NC_063383.1 | 9.73 | 0.1 | 0 | 0 | 0 |
| **MPOX** | 67 | 10244 | NC_063383.1 | 99.06 | 14.5 | 69.0 | 21.1 | 6.1 |
| **ONNV** | 22 | 2169701 | NC_001512.1 | 99.97 | 415.2 | 98.8 | 97.9 | 97.1 |

*__Supplementary Table 2__ continued*

| Name | studyID | NCBI TaxID | Chromosome | Coverage (%) | Depth of Coverage | Positions > 10 reads (%) | Positions > 20 reads (%) | Positions > 30 reads (%) |
|---|---|---|---|---|---|---|---|---|
| **RRV** | 23 | 11029 | NC_075016.1 | 100.0 | 813.1 | 99.7 | 99.6 | 99.5 |
| **RVFV** | 24 | 11588 | NC_014395.1 | 100.0 | 168.6 | 100.0 | 99.8 | 90.2 |
| **RVFV** | 24 | 11588 | NC_014396.1 | 100.0 | 270.1 | 100.0 | 99.1 | 98.6 |
| **RVFV** | 24 | 11588 | NC_014397.1 | 100.0 | 298.7 | 100.0 | 100.0 | 100.0 |
| **SARS-CoV-2** | 25 | 2697049 | NC_045512.2 | 100.0 | 400.2 | 99.9 | 99.8 | 99.8 |
| **SARS-CoV-2** | 26 | 2697049 | NC_045512.2 | 99.9 | 294.0 | 99.8 | 99.7 | 99.7 |
| **SARS-CoV-2** | 27 | 2697049 | NC_045512.2 | 100.0 | 276.0 | 99.8 | 99.7 | 99.7 |
| **SARS-CoV-2** | 28 | 2697049 | NC_045512.2 | 100.0 | 520.7 | 99.9 | 99.8 | 99.7 |
| **SARS-CoV-2** | 29 | 2697049 | NC_045512.2 | 99.9 | 165.6 | 99.6 | 99.5 | 99.5 |
| **SARS-CoV-2** | 30 | 2697049 | NC_045512.2 | 99.9 | 330.8 | 99.7 | 99.6 | 99.6 |
| **SARS-CoV-2** | 31 | 2697049 | NC_045512.2 | 99.9 | 95.2 | 99.8 | 99.5 | 99.3 |
| **SARS-CoV-2** | 32 | 2697049 | NC_045512.2 | 99.9 | 77.9 | 99.7 | 99.4 | 97.2 |
| **SARS-CoV-2** | 69 | 2697049 | NC_045512.2 | 100.0 | 388.7 | 99.8 | 99.8 | 99.8 |
| **SARS-CoV-2** | 70 | 2697049 | NC_045512.2 | 99.8 | 96.8 | 99.6 | 99.4 | 98.5 |
| **SFNV** | 35 | 206160 | NC_078077.1 | 100.0 | 184.3 | 99.8 | 98.8 | 98.1 |
| **SFNV** | 35 | 206160 | NC_078078.1 | 100.0 | 182.1 | 99.8 | 98.9 | 98.0 |
| **SFNV** | 35 | 206160 | NC_078079.1 | 100.0 | 274.2 | 99.4 | 97.3 | 95.8 |
| **SFNV** | 42 | 206160 | NC_078077.1 | 25.9 | 1.9 | 5.2 | 2.5 | 2.2 |
| **SFNV** | 42 | 206160 | NC_078078.1 | 0 | 0 | 0 | 0 | 0 |
| **SFNV** | 42 | 206160 | NC_078079.1 | 36.2 | 0.8 | 0 | 0 | 0 |
| **SFTV** | 36 | 688699 | NC_015411.1 | 99.8 | 214.4 | 96.9 | 93.4 | 92.3 |
| **SFTV** | 36 | 688699 | NC_015412.1 | 100.0 | 222.2 | 99.5 | 98.3 | 96.8 |
| **SFTV** | 36 | 688699 | NC_015413.1 | 100.0 | 573.4 | 100.0 | 100.0 | 99.8 |
| **SINV** | 37 | 11034 | NC_001547.1 | 100.0 | 411.9 | 99.8 | 99.5 | 98.5 |
| **TBEV** | 38 | 11084 | NC_001672.1 | 97.3 | 608.5 | 96.1 | 95.4 | 94.5 |
| **TBEV** | 39 | 11084 | NC_001672.1 | 97.2 | 524.0 | 95.3 | 93.9 | 93.9 |
| **TBEV** | 40 | 11084 | NC_001672.1 | 96.8 | 159.6 | 93.9 | 93.8 | 93.7 |
| **TBEV** | 41 | 11084 | NC_001672.1 | 98.4 | 132.0 | 96.8 | 96.2 | 95.4 |
| **USUV** | 43 | 64286 | NC_006551.1 | 100.0 | 467.9 | 99.4 | 99.0 | 98.9 |
| **USUV** | 44 | 64286 | NC_006551.1 | 100.0 | 867.5 | 99.8 | 99.4 | 99.0 |
| **VEEV** | 46 | 11036 | NC_075022.1 | 100.0 | 957.6 | 99.8 | 99.7 | 99.6 |
| **WEEV** | 47 | 11039 | NC_075015.1 | 98.6 | 550.2 | 98.3 | 98.3 | 98.3 |
| **WNV** | 48 | 11082 | NC_009942.1 | 100.0 | 639.2 | 99.9 | 99.3 | 99.2 |
| **WNV** | 49 | 11082 | NC_009942.1 | 100.0 | 432.8 | 99.2 | 99.1 | 99.1 |
| **WNV** | 50 | 11082 | NC_009942.1 | 60.4 | 9.28 | 23.0 | 14.3 | 11.9 |
| **WNV** | 51 | 11082 | NC_009942.1 | 77.2 | 17.6 | 42.3 | 27.1 | 18.6 |

***Supplementary Table 2** continued*

| Name | studyID | NCBI TaxID | Chromosome | Coverage (%) | Depth of Coverage | Positions > 10 reads (%) | Positions > 20 reads (%) | Positions > 30 reads (%) |
|---|---|---|---|---|---|---|---|---|
| **YFV** | 52 | 11089 | NC_002031.1 | 99.9 | 415.3 | 99.3 | 99.0 | 99.0 |
| **YFV** | 53 | 11089 | NC_002031.1 | 100.0 | 691.7 | 99.9 | 99.2 | 99.1 |
| **YFV** | 54 | 11089 | NC_002031.1 | 99.9 | 184.2 | 99.0 | 98.9 | 98.9 |
| **ZIKV** | 55 | 64320 | NC_035889.1 | 99.9 | 228.6 | 98.9 | 98.8 | 98.8 |
| **ZIKV** | 56 | 64320 | NC_035889.1 | 99.5 | 377.4 | 98.9 | 98.8 | 98.4 |
| **ZIKV** | 57 | 64320 | NC_035889.1 | 99.0 | 173.7 | 98.7 | 98.6 | 97.4 |
| **ZIKV** | 58 | 64320 | NC_035889.1 | 99.6 | 192.2 | 98.7 | 97.7 | 96.9 |

***Supplementary Table 3****: Coverage statistics for clinical Plasmodium samples in SquiDBase. For each of the three patient samples (PS1, PS2, and PS3), depth and coverage are presented relative to the Plasmodium falciparum 3D7 reference genome. Further details on the sequencing methodology can be found in De Meulenaere et al. (2024)*[13].

| Region | PS1 Mean depth | PS1 Coverage (%) | PS2 Mean depth | PS2 Coverage (%) | PS3 Mean depth | PS3 Coverage (%) | PS3 Mean depth | PS3 Coverage (%) |
|---|---|---|---|---|---|---|---|---|
| **full genome** | 19.7 | 99.8 | 5.6 | 97.4 | 351.5 | 99.9 | 99.9 | 335.1 |
| **chr 1** | 19.6 | 99.5 | 6.1 | 97.0 | 335.9 | 99.8 | 99.8 | 332.0 |
| **chr 2** | 19.9 | 99.8 | 5.2 | 97.2 | 356.6 | 100.0 | 100.0 | 344.1 |
| **chr 3** | 20.2 | 99.9 | 6.0 | 97.4 | 359.3 | 100.0 | 100.0 | 357.0 |
| **chr 4** | 19.1 | 99.5 | 5.3 | 94.7 | 371.2 | 99.9 | 99.9 | 348.9 |
| **chr 5** | 19.2 | 99.9 | 5.2 | 98.2 | 340.5 | 99.9 | 99.9 | 324.9 |
| **chr 6** | 19.7 | 99.6 | 5.5 | 96.9 | 346.6 | 99.7 | 99.8 | 340.1 |
| **chr 7** | 18.4 | 99.1 | 5.6 | 95.0 | 341.8 | 99.6 | 99.9 | 328.9 |
| **chr 8** | 19.9 | 99.9 | 5.7 | 95.7 | 334.8 | 99.7 | 99.8 | 316.3 |
| **chr 9** | 19.6 | 99.9 | 5.5 | 98.1 | 354.3 | 99.9 | 100.0 | 340.8 |
| **chr 10** | 19.4 | 99.9 | 5.4 | 97.8 | 339.6 | 100.0 | 100.0 | 323.0 |
| **chr 11** | 20.2 | 99.8 | 5.5 | 97.9 | 350.6 | 99.9 | 100.0 | 325.5 |
| **chr 12** | 19.0 | 99.8 | 5.9 | 96.8 | 344.1 | 99.9 | 100.0 | 327.8 |
| **chr 13** | 20.3 | 99.9 | 5.5 | 98.6 | 360.0 | 100.0 | 100.0 | 349.4 |
| **chr 14** | 19.6 | 99.9 | 5.7 | 98.4 | 349.6 | 100.0 | 100.0 | 332.6 |

### Supplementary Figures

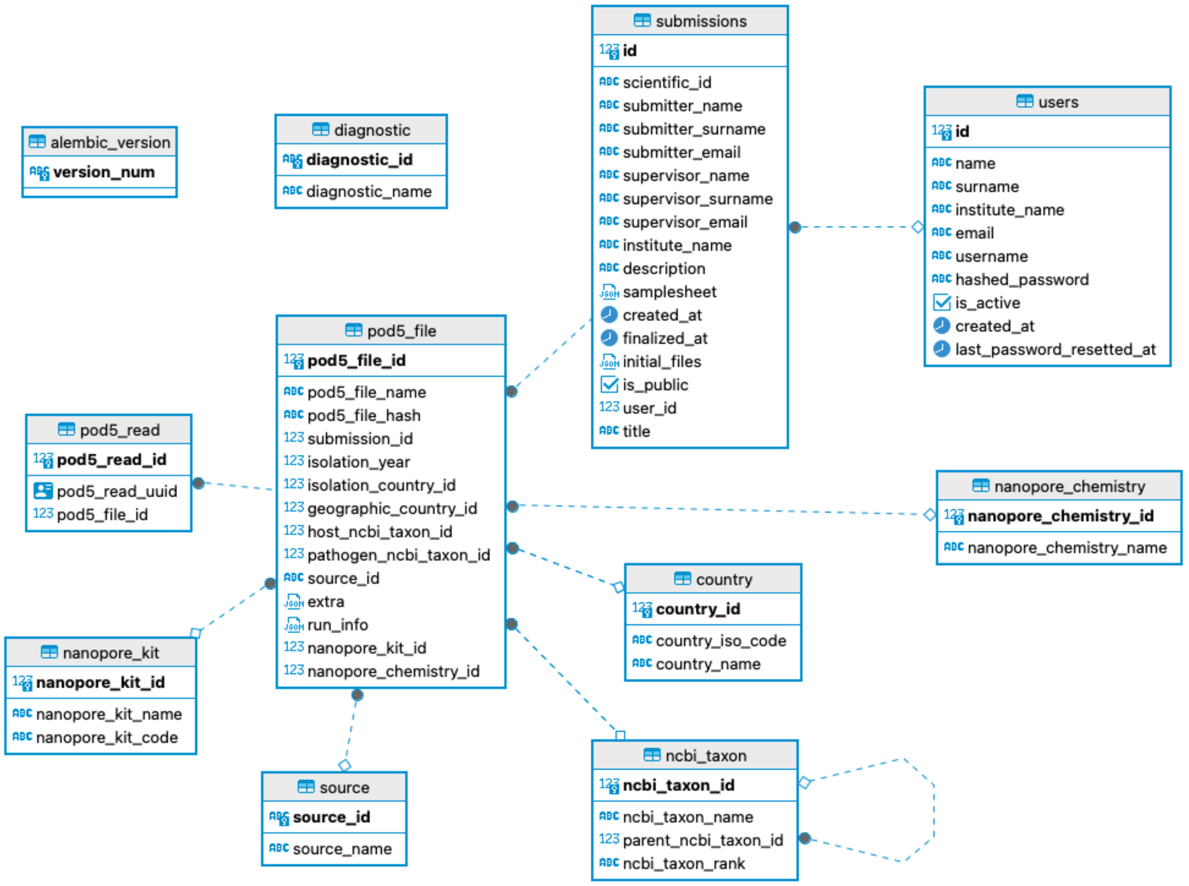

***Supplementary Figure 1**: Structure of the SQL part of SquiDBase.*

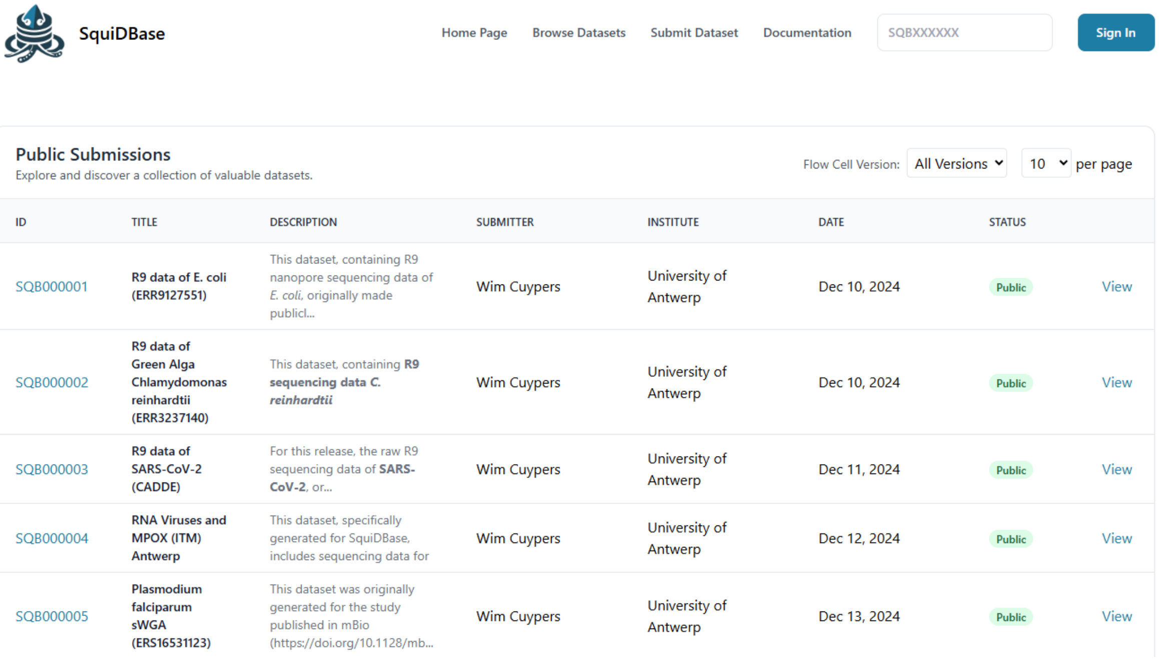

| ID | TITLE | DESCRIPTION | SUBMITTER | INSTITUTE | DATE | STATUS | |
|---|---|---|---|---|---|---|---|
| SQB000001 | **R9 data of E. coli (ERR9127551)** | This dataset, containing R9 nanopore sequencing data of *E. coli*, originally made publicl... | Wim Cuypers | University of Antwerp | Dec 10, 2024 | Public | View |
| SQB000002 | **R9 data of Green Alga Chlamydomonas reinhardtii (ERR3237140)** | This dataset, containing **R9 sequencing data *C. reinhardtii*** | Wim Cuypers | University of Antwerp | Dec 10, 2024 | Public | View |
| SQB000003 | **R9 data of SARS-CoV-2 (CADDE)** | For this release, the raw R9 sequencing data of **SARS-CoV-2**, or... | Wim Cuypers | University of Antwerp | Dec 11, 2024 | Public | View |
| SQB000004 | **RNA Viruses and MPOX (ITM) Antwerp** | This dataset, specifically generated for SquiDBase, includes sequencing data for | Wim Cuypers | University of Antwerp | Dec 12, 2024 | Public | View |
| SQB000005 | **Plasmodium falciparum sWGA (ERS16531123)** | This dataset was originally generated for the study published in mBio (https://doi.org/10.1128/mb... | Wim Cuypers | University of Antwerp | Dec 13, 2024 | Public | View |

***Supplementary Figure 2**: The 'Browse Datasets' page in SquiDBase displays each dataset with a unique SQB identifier, title, description, and details about the submitter (name, institute) along with the submission date. This page is accessible at https://squidbase.org/submissions.*

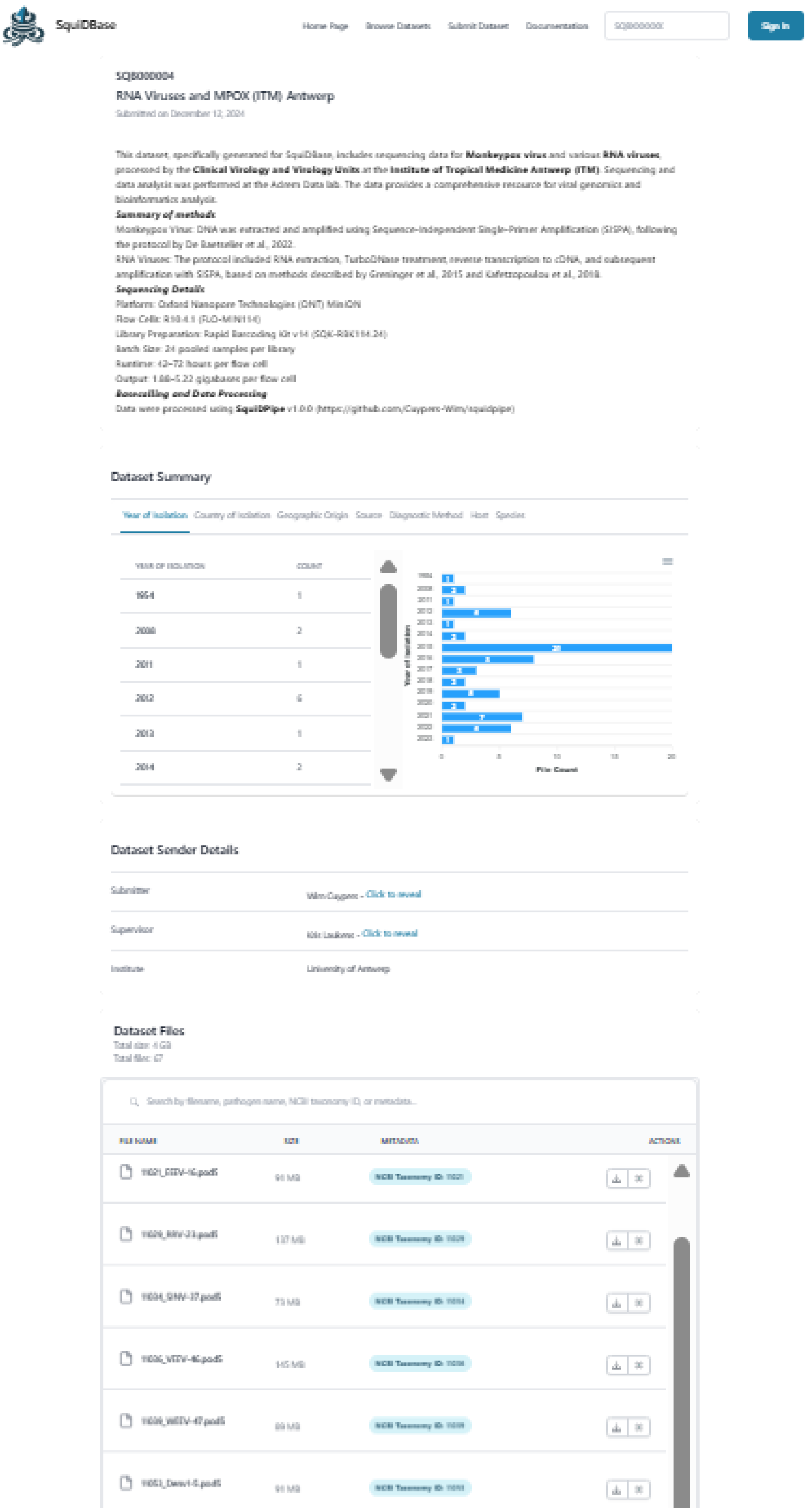

***Supplementary Figure 3**: Dataset entry linked to a SquiDBase identifier. The page shows a textual summary of the dataset, followed by a visual summary of the available metadata followed by a 'Dataset Files' screen to download all data files separately. This screenshot was generated from https://squidbase.org/submissions/SQB000004.*

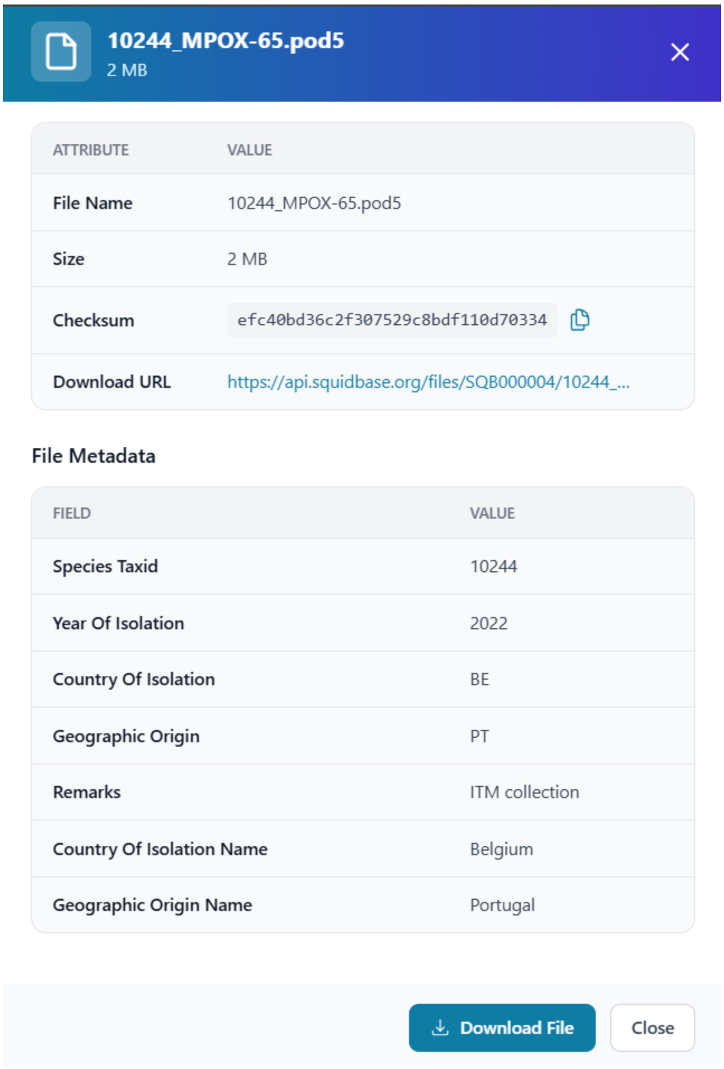

***Supplementary Figure 4***: *In the 'Dataset Files' section for each dataset submission, the eye icon expands to reveal complete metadata, including an MD5 checksum for file integrity verification.*
